# Transcription factor SmWRKY1 positively promote the biosynthesis of tanshinones in *Salvia miltiorrhiza*

**DOI:** 10.1101/230029

**Authors:** Wenzhi Cao, Yao Wang, Min Shi, Xiaolong Hao, Weiwei Zhao, Yu Wang, Jie Ren, Guoyin Kai

## Abstract

Tanshinones, one group of bioactive diterpenes, were widely used in the treatment of cardiovascular diseases. WRKYs play important roles in plant metabolism, but their regulation mechanism in S. miltiorrhiza remains elusive. In this study, one WRKY transcription factor SmWRKY1 was isolated and characterized from S. miltiorrhiza. Multiple sequence alignment and phylogenetic tree analysis showed SmWRKY1 shared high homology with other plant WRKYs such as CrWRKY1. SmWRKY1 were predominantly expressed in leaves and stems, and was responsive to salicylic acid (SA), methyl jasmonate (MeJA) and nitric oxide (NO) treatment. Subcellular localization analysis found that SmWRKY1 was localized in the nucleus. Over-expression of SmWRKY1 significantly elevated the transcripts of genes involved in MEP pathway especially 1-deoxy-D-xylulose 5-phosphate synthase (SmDXS) and 1-deoxy-D-xylulose 5-phosphate reductoisomerase (SmDXR), resulted in over 6 folds increase in tanshinones production in transgenic lines (up to 13.731mg/g dry weight (DW)) compared with the control lines. Dual-luciferase (Dual-LUC) assay showed that SmWRKY1 can positively regulate SmDXR expression by binding to its promoter. Our work revealed that SmWRKY1 participated in the regulation of tanshinones biosynthesis and acted as a positive regulator through activating SmDXR in the MEP pathway, thus discloses a new insight to further excavate the regulation mechanism of tanshinones biosynthesis.

## Introduction

*Salvia miltiorrhiza* Bunge, belonging to the *Lamiaceae* family, is a famous and prevalent Chinese herbal plant that has been widely used for the treatment of cardiovascular and cerebrovascular diseases (Zhang et al., 2010; Kai et al., 2011). The abietane-type diterpenes in *S. miltiorrhiza* are the liposoluble tanshinones including dihydrotanshinone, tanshinone I, tanshinone IIA and cryptotanshinone, which exert a variety of biological activities such as antioxidant, heart-protection, antibacterial and antitumor (Zhang et al., 2011; Chen et al., 2012; Gong et al., 2012; Xu et al., 2010, 2015). However, serious quality decrease and the low content of tanshinones in cultivated *S. miltiorrhiza* greatly limited the increasing market need (Hao et al. 2015; Zhou et al., 2016a). Therefore, it is important to improve the content of tanshinones by genetic engineering, which relies on deep understanding of the tanshinone biosynthetic pathway to *S. miltiorrhiza* (Liao et al., 2009; Zhou et al., 2016a). Tanshinones derived from the terpenoids metabolism including the mevalonate (MVA) pathway and the 2-C-methyl-d-erythritol-4-phosphate (MEP) pathway, both consisted of a series of complex enzyme catalytic reactions while operated in separate subcellular compartments, the MVA pathway localized in the cytosol and the MEP pathway took place in plastids (Kai et al., 2011; Shi et al., 2016a). Isopentenyl pyrophosphate (IPP) and its isomer dimethylallyl pyrophosphate (DMAPP) from the MEP pathway are the universal C5 precursors of tanshinones, therefore, tanshinone are generally considered to be mainly derived from the MEP pathway (Ge and Wu, 2005; Yan et al., 2009; Kai et al., 2011; Zhou et al., 2016). Recently, several key genes including 3-hydroxy-3-methylglutaryl CoA reductase (HMGR), 1-deoxy-D-xylulose-5-phosphate reductoisomerase (DXR), 1-deoxy-D-xylulose-5-phosphate synthase (DXS), geranylgeranyl diphosphate synthase (GGPPS), copalyl diphosphate synthase (CPS), kaurene synthase (KS), miltiradiene oxidase (CYP76AH1) have been successfully cloned and characterized from *S.miltiorrhiza* (Liao et al., 2009; Yan et al., 2009; Kai et al., 2010; Ma et al., 2012; Shi et al., 2014; Gao et al., 2009; Guo et al, 2013; Zhou et al., 2016a). Clarification of the above key genes involved in the tanshinones biosynthetic pathway enabled us to produce elevated concentration of tanshinones in *S. miltiorrhiza* through genetic engineering by manipulating one or several regulation points in either MVA or MEP pathway (Kai et al., 2011; Shi et al., 2014; Shi et al., 2016a). Co-expression of *SmHMGR* and *SmGGPPS* increased tanshinone production significantly in a transgenic *S. miltiorrhiza* hairy root line HG9 (Kai et al., 2011). In addition, introduction of *SmHMGR* and *SmDXR* into *S. miltiorrhiza* hairy roots enhanced the content of tanshinones apparently (Shi et al., 2014). Simultaneous introduction of *SmGGPPS* and *SmDXSII* into *S. miltiorrhiza* hairy root significantly improved the production of tanshinones, besides, their expression in *Arabidopsis thaliana* plants increased production of carotenoids, gibberellins and chlorophyll in contrast to the non-transgenic lines (Shi et al., 2016a).

Apart from manipulation of pivotal catalytic steps in the biosynthetic process of tanshinones, regulation of transcription factors such as MYB and bHLH transcription factor family was also considered as a feasible strategy to mine the biosynthesis mechanism of tanshinones (Zhou et al., 2016; Zhang et al., 2017). Heterologous expression of maize transcription factor C1 in *S.miltiorrhiza* hairy roots elevated the accumulation of tanshinones through direct interaction of C1 with its recognition sequences of pathway genes, especially mevalonate-5-diphosphate-decarboxylase (*SmMDC*) and 5-phosphomevalonate kinase (*SmPMK*) to upregulate their expression levels (Zhao et al., 2015). RNA interference (RNAi) of *SmMYC2a/b* affected multiple genes in tanshinone biosynthetic pathway and led to a reduction of tanshinones contents, implying that *SmMYC2a/b* may be a positive regulator of tanshinones accumulation (Zhou et al., 2016b). Overexpression of a *S. miltiorrhiza* R2R3-MYB gene *SmMYB9b* resulted in a 2.2-fold enhancement of tanshinones accumulation in danshen hairy roots over the control (Zhang et al., 2017). However, less is known about WRKY transcription factors and their regulation mechanism in *S. miltiorrhiza* (Li et al., 2015).

Salicylic acid (SA) is a kind of import plant hormone signal in plant metabolism, which is also reported that it could induce the accumulation of tanshionone as reported before (Hao et al., 2015), but its regulation mechanism is not clear yet. WRKY transcription factors form one of the largest gene families unique to plants, which are involved in plant secondary metabolism(Suttipanta et al., 2011). The first WRKY gene named *SPF1*, was identified from sweet potato (Ishiguro and Nakamura, 1994). Subsequently, much attention has been paid to identify and analyze WRKY genes from different model and crop plants, for instance *Arabidopsis* (Eulgem et al., 2000; Kalde et al., 2003; Wang et al., 2011), soybean (*Glycine max*) (Yin et al., 2013), *tobacco* (Yoda et al., 2002), rice (*Oryza sativa*) (Wu et al., 2005) and so on. Meanwhile, WRKY has been isolated from some traditional herbal plants including *Artemisia annua, Coptis japonica* and *Catharanthus roseus* (Jiang et al., 2015; Chen et al., 2017; Kato et al., 2007; Suttipanta et al., 2011). The significant feature of WRKY transcription factor is their WRKY domain which is approximately 60-amino acid long with the highly conserved amino acid sequence WRKYGQK located at the N-terminal and a non-typical zinc-finger-like motif C2HC (C–X_7_–C–X_23_–H–X_1_–C) or C2H2(C–X_4–5_–C–X_22–23_–H–X_1_–H) at the C-terminus (Xu et al., 2004; Lu et al., 2015). WRKY proteins can bind to the W-box cis-elements (T)TGAC(C/T) in the promoter region of some defense-related genes (Xu et al., 2004; Rushton et al., 2010; Liu et al., 2016). WRKY transcription factors can be separated into three sub-groups in accordance with the number of specific WRKY domains and zinc-finger-like motifs, Group I contains two WRKY domains and C2H2 motif, Groups II has one WRKY domain and C2H2 motif and Group III possesses one WRKY domain and C2HC motif (Eulgem et al., 2000; Rushton et al., 2010). WRKYs have shown many different functions on multiple physiology activities including stress defense, trichome development and secondary metabolism (Jiang et al., 2016). For example, *Gossypium arboretum* WRKY1 (GaWRKY1) was found to participate in regulation of sesquiterpene biosynthesis in cotton by regulate the target gene (+)-delta-cadinene synthase (CAD1) (Xu et al., 2004). *C. roseus* WRKY1 (CrWRKY1) bound to the W-box elements of the tryptophan decarboxylase (TDC) promoter involved in terpenoid indole alkaloid (TIA) biosynthetic pathway and accumulated up to 3-fold higher levels of serpentine compared with control hairy roots (Suttipanta et al., 2011). The WRKY transcription factor *GLANDULAR TRICHOME-SPECIFIC WRKY 1* (*AaGSW1*) positively regulated the expression of *AaCYP71AV1* and *AaORA* by conjunction to the W-box motifs in their promoters (Chen et al., 2017). *Glycine max* WRKY27 responsive to various abiotic stresses interacted with GmMYB174, and then cooperatively inhibited *GmNAC29* expression, facilitating stress-tolerance of drought and cold in soybean (Wang et al., 2015). A WRKY transcription factor from *W. somnifera* bound to the W-box region in the promoters of squalene synthase and squalene epoxidase, regulating the accumulation of triterpenoids in *W. somnifera* including phytosterols and withanolides (Singh et al., 2017). However, functional WRKYs related to secondary metabolism of tanshinones or salvianolic acids in *S. miltiorrhiza* have not been reported.

In this study, a WRKY transcription factor has been isolated from *S. miltiorrhiza* (named as *SmWRKY1*) and functionally characterized. Phylogenetic analysis showed that it shared high homology with AtWRKY70, CrWRKY1 and GaWRKY1. Multiple sequence alignment revealed that the nucleus-localized *SmWRKY1* contained one WRKY domain, with conserved amino acid sequence WRKYGQK and a C2HC type zinc-finger-like motif, therefore it can be classified into group III WRKY transcription factors. Introduction of *SmWRKY1* into *S. miltiorrhiza* hairy roots improved the transcripts of *SmDXS* and *SmDXR* involved in MEP pathway, resulting in higher level of tanshinones in transgenic lines compared with the control lines (2.175mg/g DW). The highest content of tanshinones was detected in *SmWRKY1-3* at 13.731mg/g DW, which was 5.3 folds higher than the control. Dual-LUC assay revealed that *SmWRKY1* activated the expression of *SmDXR* by binding to the promotor region containing one w-box in vivo. Taken together, our work revealed that *SmWRKY1* positively elevated the accumulation of tanshinones, which provides a new insight to further excavate the regulation mechanism of tanshinones biosynthesis.

## Materials and Methods

### Plant samples and reagents

*S. miltiorrhiza* seedlings used for Agrobacterium-mediated transformation were cultivated in Murashige and Skoog (MS) medium (pH5.8) containing 3% sugar and 0.8% agar in the greenhouse, growth conditions were as follows: 16 h: 8 h, light: dark cycle under 25°C with 60% relative air humidity as reported before (Kai et al., 2011; Shi et al., 2014, 2016a). Seeds of N. benthamiana were sown and cultivated in the pots supplemented with soil matrix for 4-5 weeks for infiltration (Zhou et al., 2016a).

All strains (*Escherichia coli* DH5α, Agrobacterium C58C1, GV3101 and ASE) and plasmid vectors (*pCAMBIA2300*, *pMON530*) used in this paper were preserved in our laboratory. The intermediate cloning vector pMD-18T and reverse transcriptase M-MLV were purchased from TaKaRa Biotechnology Co., Ltd. Primers-synthesizing and DNA sequencing were performed by Shanghai Sangon Biotechnological Company, China. RNA extraction kit and qRT-PCR kit were purchased from Tiangen Company. Standards of cryptotanshinone, tanshinone I, tanshinone IIA, dihydrotanshinone used for HPLC analysis were purchased from Aladdin, China. MJ, SA and SNP used for elicitation treatments were purchased from Sigma-Aldrich, Sinopharm Chemical Reagent Co., Ltd, respectively.

### Elicitor preparation

For methyl jasmonate (MeJA) induction, MeJA was dissolved in 5% ethanol, and then dissolved into distilled water to a storage concentration of 50 mM. A final working concentration of 100 μM MeJA was employed for elicitation assay, and equivalent volume of sterilized water was used as the mock treatment (Kai et al., 2012). For salicylic acid (SA) treatment, SA was dissolved in sterile water to a storage concentration of 50 mM, and then added to hairy roots cultures to the final concentration of 100 μM (Hao et al., 2015). For NO elicitation, first a concentration of 100 mM SNP solution was obtained, and then applied to cultures to 100 μM. All the above-mentioned solutions were sterilized through 0.22μm filters (Pall Corporation, USA). And solvent of the equivalent volume was added into the control group.

### Identification and cloning of *SmWRKY1*

A local transcription database of *S. miltiorrhiza* built up as reported previously (Shi et al., 2016b) was used for this research. One partial *WRKY* in high homology with other plants *WRKYs* while lack of 3’-terminal was chosen for further study. Gene-specific forward primer *SmWRKY1*-F605 was designed to amplify the 3’ end of *SmWRKY1* as well as the reverse primer AUAP by rapid amplification of cDNA ends (RACE) (Liao et al., 2009; Kai et al., 2010; Zhang et al., 2011). 5’-sequence and 3’-terminal products was aligned and assembled to obtain the full-length cDNA sequence of the putative *SmWRKY1* gene. Primer pairs *SmWRKY1-KF* and *SmWRKY1-KR* were synthesized for amplification of the full ORF of *SmWRKY1* according to the procedure as described below: initial denaturation at 94 °C for 10 min, 35 cycles of 94 °C for 45 s, 55 °C for 45 s and 72 °C for 90 s, followed by a final extension at 72 °C for 10 min. All primers used for identification of *SmWRKY1* were listed in Supplemental Table 1.

### Bioinformatics analysis of *SmWRKY1*

Biological characteristics of *SmWRKY1* were further analyzed by a series of tools. Nucleotide blast, protein blast and ORF Finder were used to analyze nucleotide sequence and complete open reading frame. MEGA 6 was applied to construct a phylogenetic tree by the neighbor-joining (NJ) method and 1000 replications were performed for bootstrap values. Multiple sequences alignment between *SmWRKY1* and other plant *WRKYs* were carried out using Clustal X with default parameters (Shi et al., 2016b; Zhou et al., 2016a).

### Expression pattern of *SmWRKY1* in different tissues and under various elicitors treatments

Different tissues including taproot, stem, leaf, flower and seed were gathered from two-year-old *S. miltiorrhiza* plants in mature. Elicitor treatments were conducted on *S. miltiorrhiza* hairy roots sub-cultured for 60 days infected with *Agrobacterium* C58C1. Hairy roots were harvested at selected time points (0h, 0.5h, 1h, 2h and 4h) after MJ treatment. And for SA and NO induction, hairy roots were collected at 0h, 3h, 4h, 6h, 9h, 12h after treatment. All the treated samples were immediately frozen in liquid nitrogen and stored for analyzing the expression profiles of *SmWRKY1*.

### Subcellular localization of *SmWRKY1*

To analyze the subcellular localization of *SmWRKY1*, PCR products of *SmWRKY1* ORF with *BglII* and *KpnI* restriction sites were digested with *BglII* and *KpnI* and cloned into the vector *pMON530* to generate the vector *pMON530-SmWRKY1-GFP.* The constructed expression vector was transferred into *Agrobacterium* strain ASE and injected into forty-day-old tobacco leaves. GFP fluorescence was observed after 48h cultivation using the confocal microscope (Carl Zeiss) (Shi et al., 2016b; Zhou et al., 2016a).

### Generation of transgenic *SmWRKY1* hairy roots

The full-length coding sequence of *SmWRKY1* with restriction sites *Spe* I and *BstEII* was cloned and inserted into modified *pCAMBIA2300^sm^* vector (replace the small fragment digested by EcoR I and Hind III with the corresponding *GFP-GUSA* gene expression cassette from *pCAMBIA1304*) under the control of the CaMV 35S promoter to generate *pCAMBIA2300^sm^-SmWRKY1* as described before (Shi et al., 2016b). *A. rhizogenes* strain C58C1 containing *pCAMBIA2300^sm^-SmWRKY1* was used to infect the aseptic explants and the empty *pCAMBIA2300^sm^* was regarded as the control. The transformation procedure was the same as our previous study (Kai et al., 2011; Shi et al., 2014, 2016a, 2016b; Zhou et al., 2016a). Hairy roots in good state were sub-cultured and primer pairs *CaMV35S-F23* and *SmWRKY1-QR* were used to identify the positive colony by polymerase chain reaction (PCR) analysis, meanwhile *rolB* gene in C58C1 was detected. Genomic DNA was isolated from individual hairy root sample by the cetyltrimethyl ammonium bromide method as previously reported (Zhou et al., 2016a, c). Identified positive-colonies were segmented approximate 4 cm long for shake-flask culture in 100 mL 1/2MS medium and cultured at 25°C on an orbital shaker shaking at the speed of 100 rpm in darkness (Shi et al., 2016a, 2016b). Primers sequences were listed in Supplemental Table 1.

### Total RNA isolation and relative expression analysis via qRT-PCR

Expression profiles of *SmWRKY1* and several key enzyme genes involved in tanshinones biosynthetic pathway were investigated by real-time quantitative PCR analysis (qRT-PCR). Total RNA was extracted from different tissues with the RNA prep pure plant kit as described before (Shi et al., 2016b). Total RNA served as the template for reverse transcription (RT) reaction, the reaction conditions were according to our previous study (Shi et al., 2016b). Gene-specific primers (listed in Supplemental Table 1) for qRT-PCR were designed and analyzed the relative expression level compared with the internal reference gene *SmActin* using the relative quantitative analysis method (2-^ΔΔCT^). Amplifications were performed according to the manufacturer’s instructions: one cycle of denaturation at 95 °C for 10 min, then 40 cycles of 15 s denaturation at 95 °C, 30 s annealing at 60 °C and 30 s extension at 72 °C.

### Dual-Luciferase (Dual-LUC) assay

For the dual-luciferase (Dual-LUC) assay, the promoters of *SmDXR* and *SmDXS2* with *KpnI* and *XhoI* restriction sites were cloned into pGREEN 0800 to drive the luciferase reporters, respectively. And the complete ORF of *SmWRKY1* was inserted into the *pCAMBIA2300^sm^* vector as effector. The *pCAMBIA2300^sm^-SmWRKY1* and *pCAMBIA2300^sm^* empty plasmid were transferred into *Agrobacterium tumefaciens* strain GV3101 individually. The *pGREEN-pSmDXR*, *pGREEN-pSmDXS2* was each co-transformed with the helper plasmid pSoup19 into GV3101, and the assay was conducted as described before (Zhang et al., 2015). The *pCAMBIA2300^sm^* empty plasmid was used as a negative control. The 35S promoter-driven Renilla was taken as an internal control. Each sample were measured for three biological times. The reporter strain with effector strain was mixed with ratio of one-to-one to inject the tobacco leaves. After two days’ injection, the samples were collected for dual-LUC assay by reaction reagents according to the manufacturer (Promega).

### Tanshinones analysiss

The 60-day-old hairy roots were dried at 50 °C to constant weight in an oven. Approximate 200 mg dried hairy roots were ground into powder and immersed in 16 mL methanol/dichloromethane (3:1, v/v) for tanshinones extraction. Tanshinones extraction was carried as reported before (Hao et al., 2015). HPLC analysis was performed on Agilent 1260 apparatus equipped with a Waters reversed-phase C18 symmetry column, and the detection conditions were performed following the methods described previously (Shi et al., 2016b).

## Results

### Isolation and molecular cloning of *SmWRKY1*

WRKY transcription factor is a large family in plants which has been proven to be involved in the regulation of many physiological processes in plants including secondary metabolism (Xu et al., 2004; Suttipanta et al., 2011). By searching our local transcriptome database, a *WKRY* fragment with 5' untranslated region (UTR) but lack of partial of 3’ terminal sequence was chosen for further research because it showed high homology with *GaWRKY1* and *CrWRKY1* as well as *Arabidopsis thaliana WKRY70.* 3’ RACE technology was used to obtain a 432 bp sequence of 3’ end of the fragment. After sequence assembly, the full-length gene was cloned and designated it as *SmWRKY1*. *SmWRKY1* sequence consists of 17 bp 5'UTR, a complete 789 bp open reading frame which encodes 262 amino acids, along with 238 bp 3' UTR.

### Bioinformatics analysis of *SmWRKY1*

To further figure out the biological characteristics and phylogenetic relationship of *SmWRKY1,* a series of bioinformatics analysis were performed. Multiple alignment of SmWRKY1 with related WRKY proteins from other plant species revealed that SmWRKY1, AaWRKY1 and CrWRKY1 all contained a conserved WRKY domain (WRKYGQK) and a special zinc finger like motif in its C-terminal which falls into the group III of WRKY family (**Fig. 1A**) and indicated that they might have similar function. Then alignment of SmWRKY1 and other plant WRKYs was performed at amino acid level and a neighbor-joining tree was constructed, as shown in (Fig. 1 B).The results revealed that SmWRKY1 shared 62%, 49%, 37%, 29% identities with EgWRKY70, NtWRKY70, CrWRKY1 and AaWRKY1, respectively.

**Figure 1.**
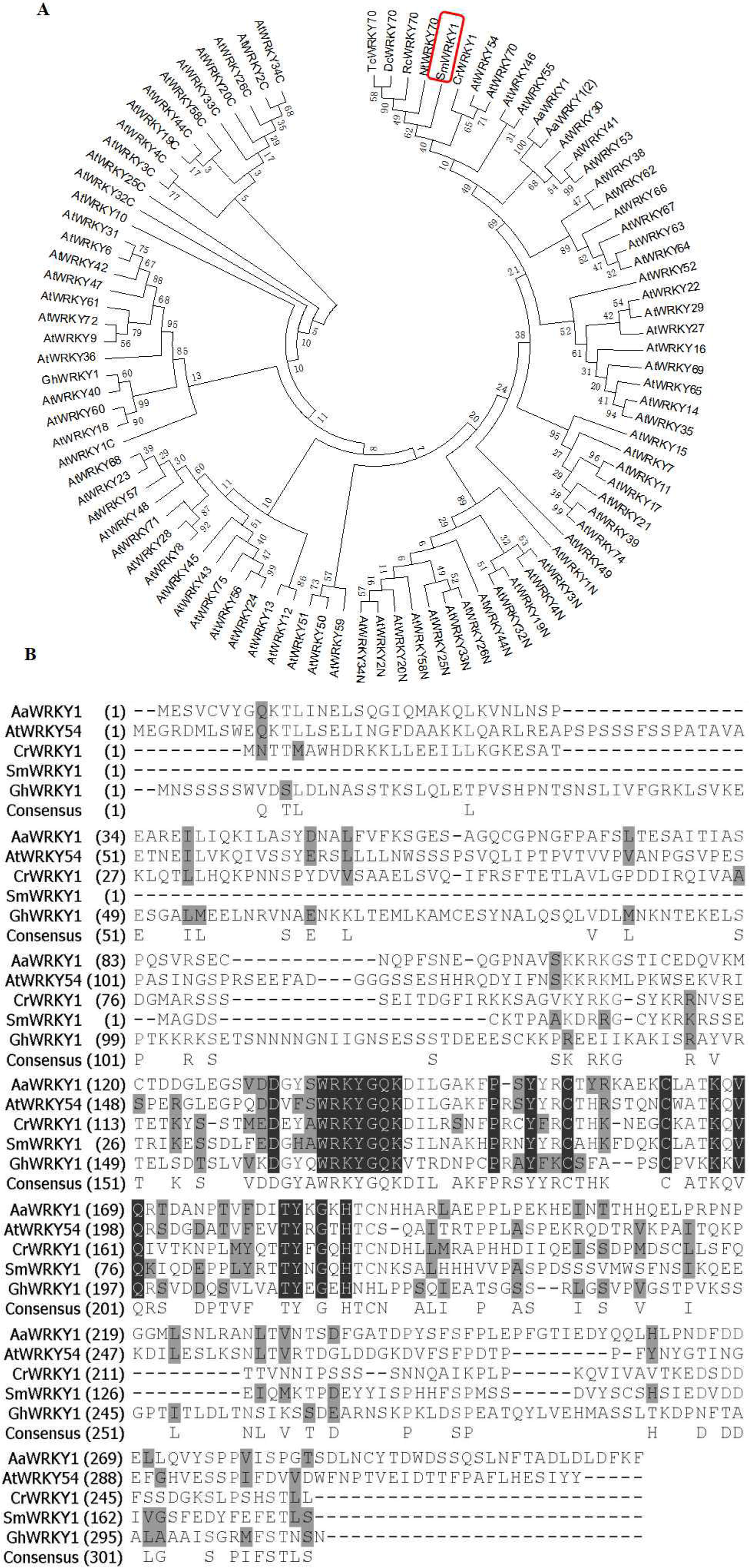
(A) Multiple alignment of *SmWRKY1* with related WRKY proteins from other plant species. Black boxes indicate identical residues; grey boxes indicate identical residues for at least three of the sequences. (**B)** Phylogenic tree analysis of SmWRKY1 and WRKY TFs from *Arabidopsis thaliana*, *Artemisia annua*, *Catharanthus roseus*, *Nicotiana tabacum*, etc. Phylogenic tree was constructed on MEGA6.0 by using neighbor-joining method and the bootstrap values were obtained for 1000 replications.

### Tissue and induction expression profiles of *SmWRKY1*

To investigate the tissue expression pattern of *SmWRKY1*, roots, stems, leaves, flowers and seeds from two-year-old *S. miltiorrhiza* plants were analyzed. *SmWRKY1* showed significant expression in leaves and stems and low expression in flower and root, its transcript was barely detected in seeds (**Fig. 2A**). This result indicated that *SmWRKY1* was not a tissue-constitutive expression gene.

**Figure 2.**
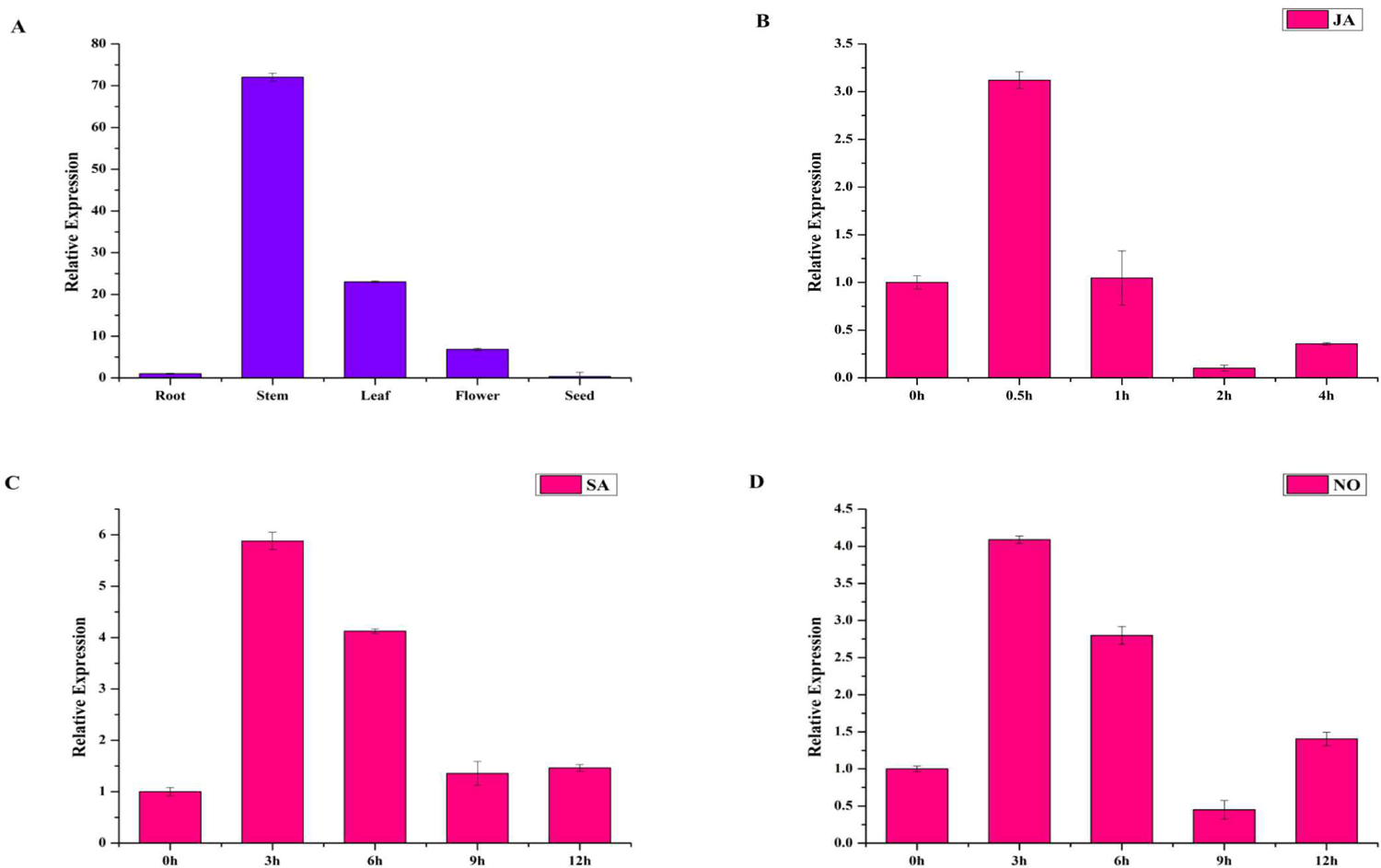
(A) Expression pattern of *SmWRKY1* in different tissues. Each tissue was obtained from several individual two-year-old *S. miltiorrhiza* plants in nature. Transcript abundance of *SmWRKY1* is normalized to actin by the method of 2^−_ΔΔ_Ct^. (**B**) The expression level of *SmWRKY1* after MeJA treatment for different time points by qRT-PCR analysis respectively. (**C**) The expression level of *SmWRKY1* after SA treatment for different time points by qRT-PCR analysis respectively. (**D**) The expression level of *SmWRKY1* after NO treatment for selected points by qRT-PCR analysis respectively.

To study whether *SmWRKY1* could respond to exogenous hormone treatment, 60-day-old *S. miltiorrhiza* hairy roots were treated with MeJA for different time points while the 0 hr point was used as control and the expression was detected by qRT-PCR. The result indicated that *SmWRKY1* expression was induced by exogenous MeJA (**Fig. 2B**), the expression level reached peak at 0.5h after treatment, arising approximate 3-fold compared with control). Then, the transcript level of *SmWRKY1* declined rapidly in two hours. Meanwhile, the hairy roots were also treated with SA and NO. Both SA and NO could induce the expression of *SmWRKY1,* which reached the maximum level at 3h and gradually decreased till 12h after treatment (**Fig. 2C, D**). In summary, *SmWRKY1* could be induced by MeJA, SA and NO.

### Subcellular localization of *SmWRKY1*

To experimentally confirm the subcellular localization of *SmWRKY1*, *SmWRKY1* was cloned into the *pMON530* vector to fuse with green fluorescent protein (GFP) reporter gene to generate vector *pMON530-SmWRKY1-GFP.* Then, the constructed vector and the *pMON530* (used as the control) was transformed into *ASE* strain and expressed in tobacco leaves, respectively. In the leaves of control vector transformed plant, the fluorescence of GFP was detected in the cytoplasm and nucleus (**Fig. 3**). On the contrast, the fluorescent signal of *SmWRKY1-fused GFP* was only examined in nucleus. The expression pattern was consistent with the character of *SmWRKY1* as a transcription factor.

**Figure 3.**
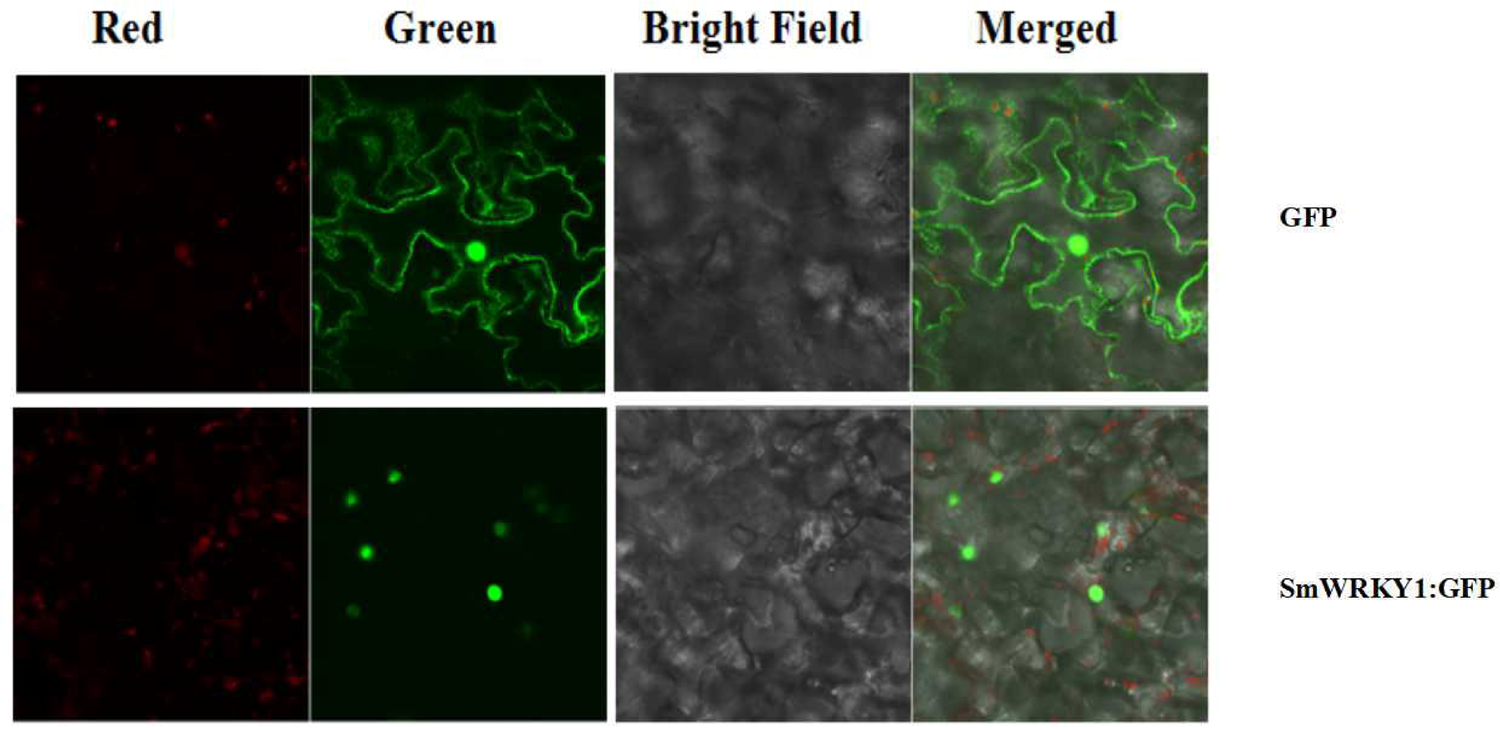
Subcellular localization of *SmWRKY1*. (**A-D**) The free GFP expressed in *N. benthamiana* leaves. (**E-H**) *SmWRKY1:: GFP* expressed in *N. benthamiana* leaves.

### Acquisition of *SmWRKY1* transgenic hairy roots

To further investigate the function of *SmWRKY1* in *S. miltiorrhiza,* we inserted *SmWRKY1* into a modified *pCAMBIA2300^sm^* vector. Then, the recombinant overexpression vector *pCAMBIA2300^sm^-SmWRKY1* was introduced into *A.rhizogenes* stain C58C1 and used to infect *S. miltiorrhiza* explants and the empty vector *pCAMBIA2300^sm^* was used as control. After 2-3 weeks the fresh hairy roots differentiated from the stem and leaf explant as shown in **Fig. 4.** The positive lines carrying *SmWRKY1* gene were verified by PCR. The positive rate was 20.5% among the 39 samples (**Fig. 5**). qRT-PCR analysis of the expression of *SmWRKY1* in over-expression lines found that *SmWRKY1* expressed 20- to 48-fold higher than the empty vector control transformed lines (**Fig. 6A).** The three high expression lines including 1, 2 and 32 (designated as 3) were chosen for further analysis.

**Figure 4.**
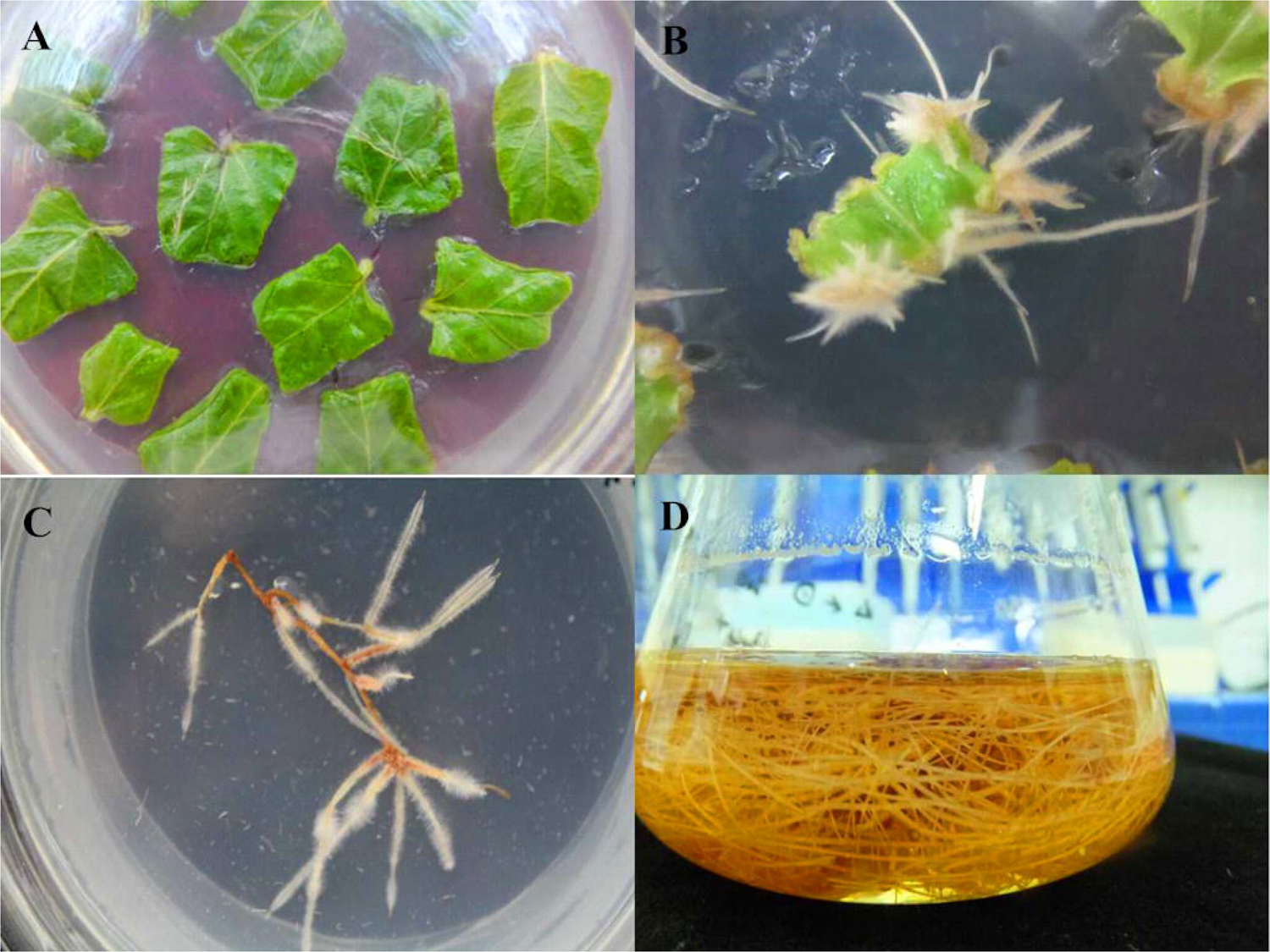
Generation of transgenic hairy root of *S. miltiorrhiza.* (**A** *S. miltiorrhiza* explants on V2MS medium; (**B**)The growing hairy root on the infected *S. miltiorrhiza* explants. (**C**) Monoclone of hairy root. (**D**) Hairy roots culture in ½MS medium.

**Figure 5.**
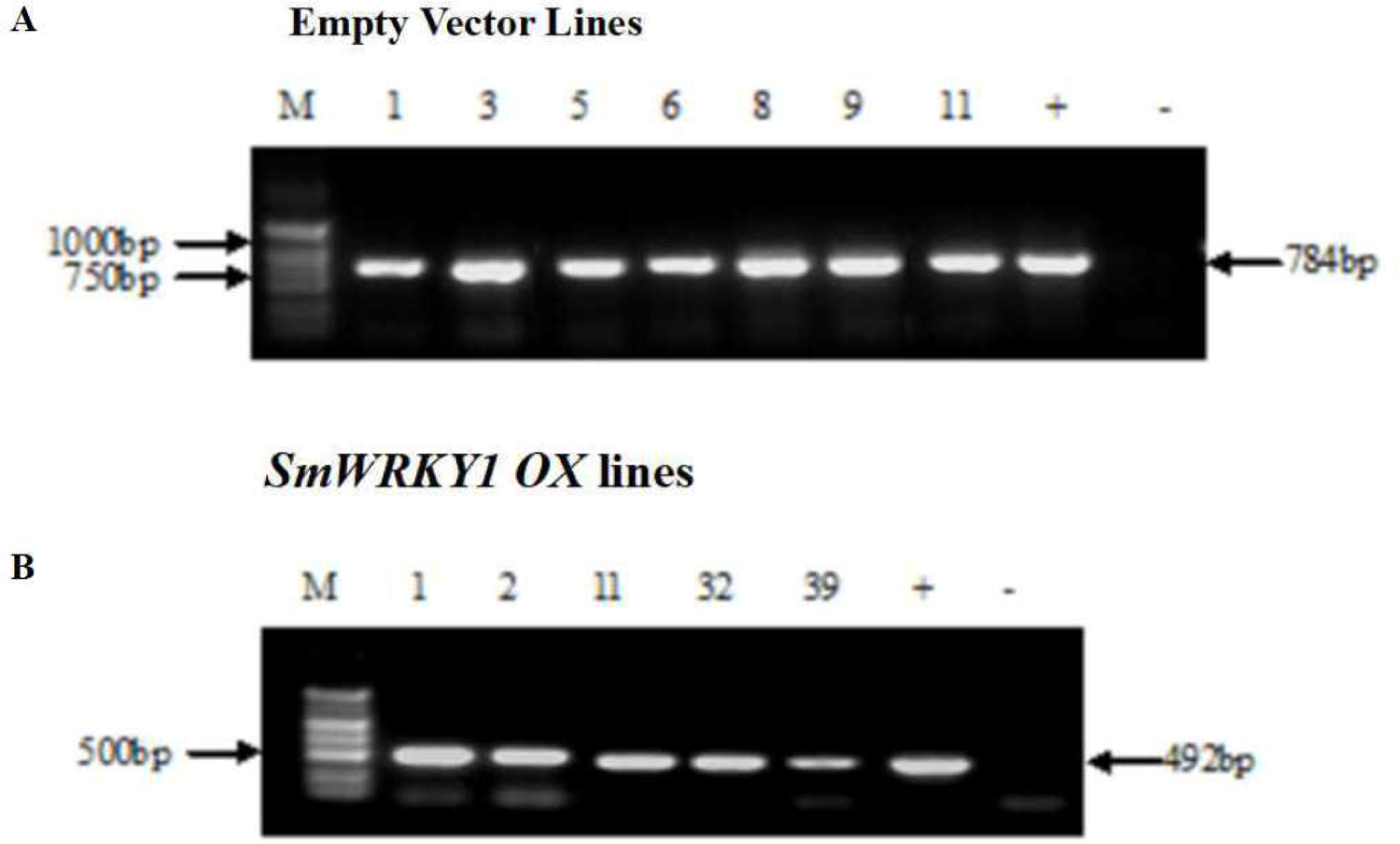
(A) Identification of positive transgenic hairy root lines by PCR. (*GusA-F* and *GusA-R* were used to identify empty vector *pCAMBIA2300^sm^* transformed lines (**B**) Primers *CaMV35S-F23* and *SmWRKY1-QR* were used to identify the positive colony of *SmWRKY1* overexpression transgenic lines).

**Figure 6.**
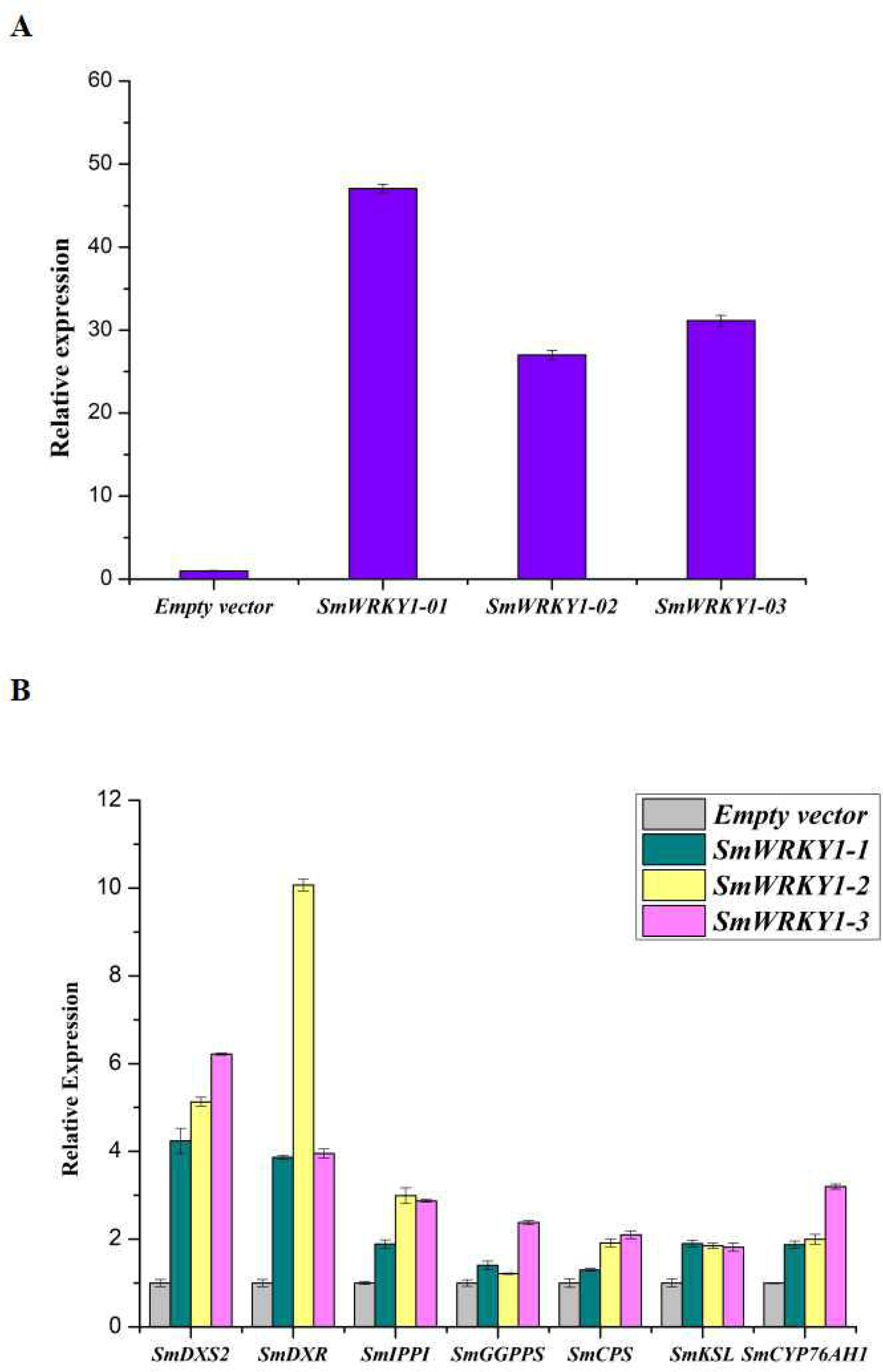
Transcript levels of *SmWRKY1* and genes related to tanshinones biosynthesis in *SmWRKY1* transgenic hairy roots. Expression of *SmWRKY1* were analyzed by qRT-PCR.

### *SmDXS* and *SmDXR* involved in MEP pathway were up-regulated by *SmWRKY1*

To study whether *SmWRKY1* participated in the regulation of tanshinone biosynthesis, transcript levels of several genes related to tanshinones biosynthesis in *SmWRKY1* transgenic hairy root were analyzed by qRT-PCR. Several tanshione biosynthesis pathway genes were up-regulated in the SmWRKY1-overexpressing hairy roots (**Fig. 6B**), the most striking ones were *SmDXS2* and *SmDXR* gene, which increased 4-6 folds and 4-10 folds compared with the control, respectively. Though the expression of *SmIPPI*, *SmGGPPS*, *SmCPS*, *SmKSL* and *SmCYP76AH1* was a little lower than *SmDXS* and *SmDXR*, their expression in over-expression lines was 2-4 folds higher than the control. In contrast, the expression of all these seven tanshinones biosynthesis pathway genes were significantly decreased in the knock-down lines. All these results suggested that *SmWRKY1* may be a positive regulator in tanshinones biosynthesis.

### SmWRKY1 activates the transcription of *SmDXR* in vivo

Expression profiles showed that *SmWRKY1* significantly promote the expression of *SmDXR* and *SmDXS2* in charge of pivotal catalytic steps of tanshinone accumulation. By analyzing the sequence of *SmDXR* and *SmDXS2* promoter,we found a W-box in the promoter of *SmDXR* (**Fig. 7A**). Than dual luciferase (dual-LUC) method was employed to verify whether SmWRKY1 protein activates the transcription of *SmDXR* and *SmDXS2* or not. The results showed that *SmWRKY1* elevated the expression of *SmDXR* by 6.08-fold(**Fig. 7B**) while endowed inconspicuous change.

**Figure 7.**
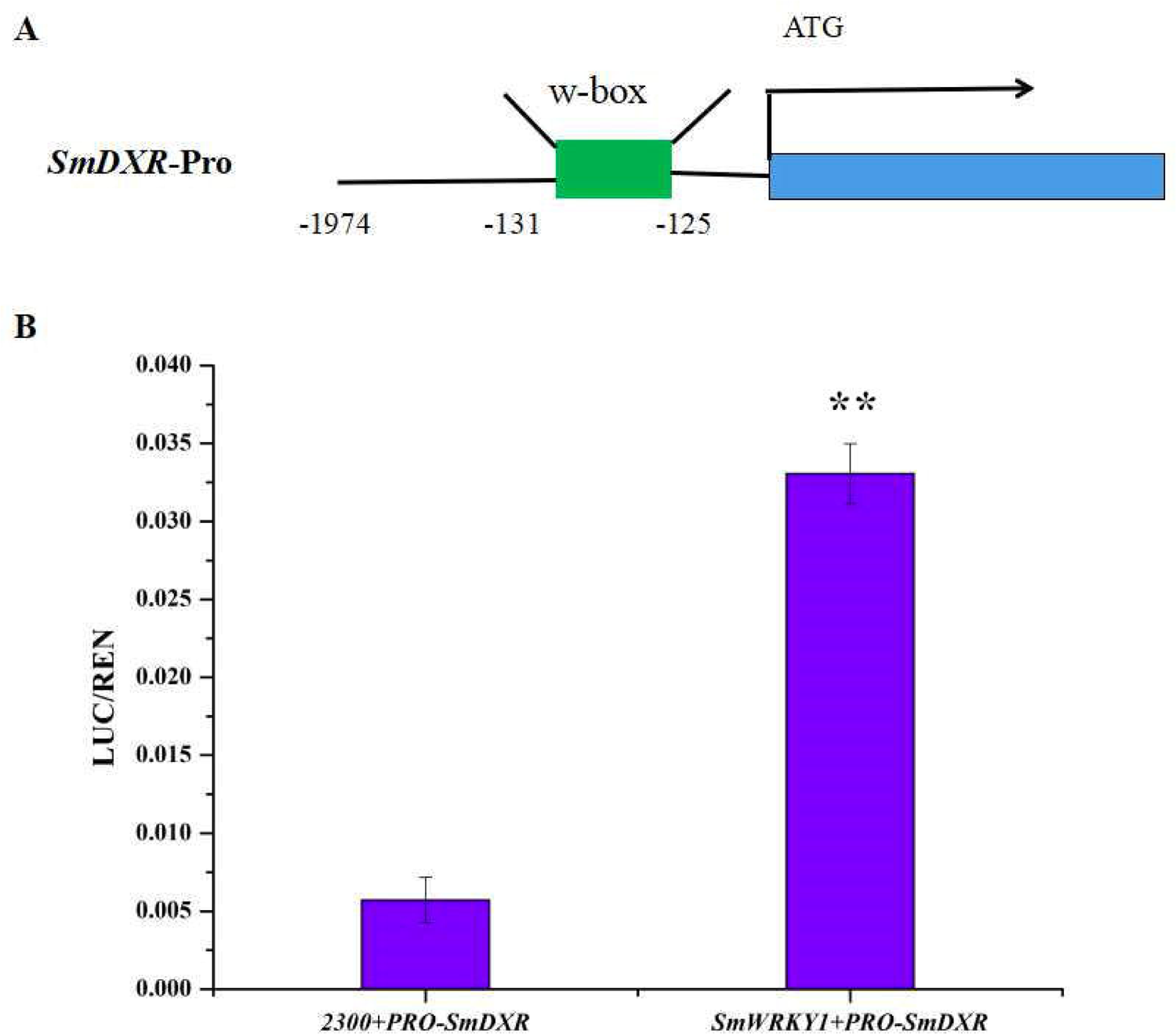
The SmDXR promoter was fused to the luciferase (LUC) reporter and the promoter activity was determined by a transient dual-LUC assay in tobacco. The value of LUC activity/ Renilla (REN) luciferase was regarded as the activating activity. Error bars indicate SD (n = 3). Student’s t-test: *, P < 0.05; **, P < 0.01.

### Accumulation of tanshinone was obviously affected by *SmWRKY1*

Based on the quantitative data, we wanted to further evaluate whether the expression of *SmWRKY1* in transgenic hairy roots affect the content of tanshinone. Three overexpression lines and two knock-down line were used to examine four monomers of tanshinone including cryptotanshinone, dihydrotanshinone I, tanshinone I, tanshinone IIA in hairy roots by HPLC. The results showed that the content of cryptotanshinone, dihydrotanshinone I, tanshinone I were significantly up-regulated and the total tanshinone had risen to 9.443-13.731mg/g DW in over expression lines. Among them *pCAMBIA2300^sm^-SmWRKY1-3* lines accumulated the highest content of total tanshinone, which was 6.31 folds higher than control (**Fig 8**). These results further confirmed the positive role of *SmWRKY1* in the regulation of tanshinone biosynthesis.

**Figure 8.**
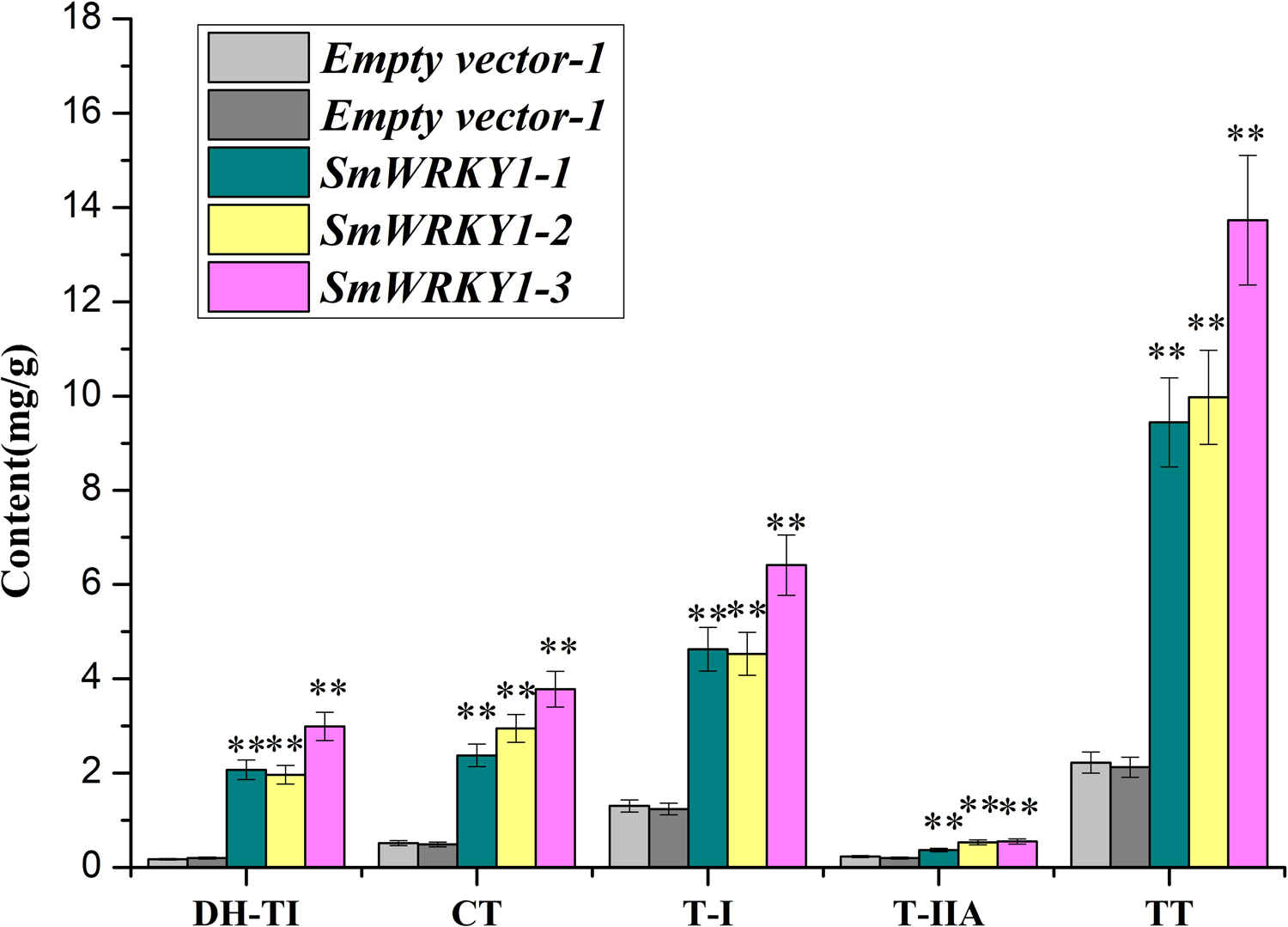
The production of tanshinone in *SmWRKY1* transgenic hairy roots compared with control detected by HPLC.

## Discussion

WRKY transcription factors are one of the largest gene families specific to plants which have been studied for decades. The conserved domain WRKYGQK and a zinc finger motif which consists of 60 amino acids are considered as the general character of WRKY TFs which also can be regarded as the criterion for subgrouping (Eulgem et al., 2000; Xie et al., 2005; Zhang and Wang, 2005). *SPF1*, *ABF1.2*, *PcWRKY1.2.3* and *ZAP1* are the first WRKY cDNAs isolated from sweet potato (*Ipomoea batatas*), wild oat (*Avena fatua*), parsley (*Petroselinum crispum*) and *Arabidopsis*, respectively (Ishiguro et al., 1994; Rushton et al., 1996; de Pater, S. et al. 1996). Up to now, 74 and 109 WRKYs members have been found in *Arabidopsis* and *Oryza sativa* respectively (Ujjal et al., 2016). Previous studies have proved that WRKY TFs could directly bind to the W-box of related genes from different signal pathways and played its regulatory role in stress tolerance in plants (Eulgem et al., 2000). For instance, *SpWRKY1* has been testified to promote resistance to *Phytophthora nicotianae* and tolerance to salt and drought stress in transgenic tobacco (Li et al., 2015). *GhWRKY25* from cotton, a member of group I, conferred transgenic *Nicotiana benthamiana* differential tolerance to abiotic and biotic stresses (Liu et al., 2016). In recent years, the role of WRKY TFs in the regulation of secondary metabolism in plants has gained attentions, and some progress has been made in this field, for example, the involvement of *Artemisia annua* WRKY1 (AaWRKY1) transcription factor can elevate the production of artemisinin by targeting the Amorpha-4,11-diene synthase (ADS) gene of *Artemisia annua* (Ma et al., 2009; Jiang et al., 2016). A jasmonate- and salicin-inducible WRKY transcription factor from *Withania somnifera* named as *WsWRKY1* could bind to W-box sequences in promoters of squalene synthase and squalene epoxidase genes in *W. somnifera* genes regarding triterpenoid biosynthesis such as phytosterol and withanolides (Singh et al., 2017). The WRKY transcription factor *GLANDULAR TRICHOME-SPECIFIC WRKY l* (*AaGSWl*) positively regulated the expression of *AaCYP7lAVl* and *AaORA* by conjunction to the W-box motifs in their promoters (Chen et al., 2017). However, lack of research on the function of WRKY TFs in *S. miltiorrhiza* especially in the regulation of tanshinone biosynthesis were reported.

*S. miltiorrhiza*, a traditional Chinese herbal medicine, has been used for thousands of years. Previous studies have proved that as a major medicinal active ingredient in *S. miltiorrhiza*, tanshinones could be used for the treatment of cardiovascular and cerebrovascular diseases in China (Chen et al., 2012). However, traditional *S. miltiorrhiza* production cannot meet the growing clinical needs due to its slow growth, low tanshinone content and scarcity of wild resources (Zhou et al., 2016a). Thus, genetic engineering has become an effective and important way to increase the accumulation of active ingredients in *S. miltiorrhiza*. Overexpression of *SmDXS* in transgenic hairy root lines can significantly enhance the production of tanshinones (Zhou et al., 2016a). Meanwhile *SmDXR* was also an important enzyme gene in tanshinone biosynthetic pathway whose overexpression could significantly improve the production of tanshinones in hairy root lines (Shi et al., 2014). In our study, a new WRKY transcription factor was successfully cloned from *S. miltiorrhiza* with high homology with *CrWRKY1* and *GaWRKY1*. qRT-PCR analysis showed that over-expression of *SmWRKY1* can promote the transcripts level of *SmDXR* and *SmDXS2* to the greatest extent in comparison to other genes involved in tanshinone biosynthetic pathway such as *SmIPPI*, *SmGGPPS*, *SmCPS*, *SmKSL* and *SmCYP76AH1*. Otherwise, dual-Luciferase (Dual-LUC) assay showed that *SmWRKY1* can positively regulate *SmDXR* expression by directly binding to the promoter region containing one W-box. HPLC analysis revealed that introduction of *SmWRKY1* in transgenic *S. miltiorrhiza* hairy roots can increase the tanshinones production up to 13.731mg/g dry weight (DW) which is over 6 folds as that in non-transgenic lines. Therefore, it is an effective strategy to regulate the tanshinone production in *S. miltiorrhiza* by introduction of related transcription factors.

To our knowledge, the defense mechanisms in plants are complicated and are mainly considered to be regulated by SA and MJ signaling network (Tsuda et al., 2009). SA plays a vital role in plant defense against pathogens and pathogen invasion obviously triggers its accumulation in plants (Qiu et al., 2009). MeJA is widely used as an elicitor to investigate the biosynthetic pathway of active compounds and the underlying regulatory mechanisms (Gundlach et al., 1992). It has been proved to be defensive to environmental stresses such as wounding, pathogen and pest attack, ozone exposure, ultraviolet radiation and salt stress as a regulator (Ma et al., 2006; Wang et al., 2011). While in *S. miltiorrhiza*, exogenous MeJA treatment can promote the accumulation of tanshinone (Gu et al., 2012; Kai et al., 2012; Hao et al., 2015). In our study, we noticed that *SmWRKY1* can be induced by exogenous MeJA treatment, reaching a maximal level at 0.5 h after MeJA treatment., which is consistent with the previous reports that MeJA treatment could increase tanshione production (Hao et al., 2015; Zhou et al., 2017). Recent studies showed that *Brassica napus* WRKY33 (BnWRKY33), a *S. sclerotiorum-responsive* gene, could positively regulate resistance to *S. sclerotiorum* by enhancing the expression of genes involved in camalexin synthesis and genes regulated by salicylic acid (SA) and jasmonic acid (JA) (Liu et al., 2017). JcWRKY a salicylic acid-inducible TF was able to work in co-ordination with SA signaling to orchestrate the different biochemical and molecular pathways to maneuvre salt stress tolerance of the transgenic tabacoo plants (Agarwal et al., 2016). Expression profiles revealed that the SmWRKY1 was responsive to both SA and MJ, which implied that SmWRKY1 may participate in the process of stress regulation such as the defense against pathogen, however need to be examined furthermore.

In conclusion, our work revealed a new transcription factor *SmWRKY1* which is involved in the regulation of tanshinone biosynthesis and promote the accumulation of tanshinone in transgenic hairy root lines by targeting *SmDXR* involved in the MEP pathway. Our study may provide a new insight by genetic engineering strategy with functional transcription factors to improve the yield of target compounds in *S. miltiorrhiza*.

## Acknowledgments

This work was supported by National Natural Science Fund (31270007, 81522049, 31571735), Zhejiang Provincial Key University Project on the Construction of First-class Subjects, New Century Talent Project (NECT-13-0902), Shanghai Science and Technology Committee Project (17JC1404300, 15430502700), the “Dawn” Program of Shanghai Education Commission (16SG38) and Shanghai Engineering Research Center of Plant Germplasm Resources (17DZ2252700).

## References

Agarwal P, Dabi M, Sapara KK, Joshi PS, Agarwal PK. 2016. Ectopic Expression of JcWRKY Transcription Factor Confers Salinity Tolerance via Salicylic Acid Signaling. Front Plant Sci 7:1541. doi:10.3389/fpls.2016.01541.

Chen J, Shi DY, Liu SL, Zhong L. 2012. Tanshinone IIA induces growth inhibition and apoptosis in gastric cancer in vitro and in vivo. Oncol Rep 27 (2):523–528. doi:10.3 892/or.2011.1524.

Chen M, Yan T, Shen Q, et al. 2017. GLANDULAR TRICHOME-SPECIFIC WRKY 1 promotes artemisinin biosynthesis in *Artemisia annua*. New Phytol 214 (1):304–316. doi:10.1111/nph.14373.

de Pater S, Greco V, Pham K, Memelink J, Kijne J. 1996. Characterization of a zinc-dependent transcriptional activator from *Arabidopsis*. Nucleic Acids Res 24 (23):4624–4631.

Eulgem T, Rushton PJ, Robatzek S, Somssich IE. 2000. The WRKY superfamily of plant transcription factors. Trends Plant Sci 5 (5):199–206.

Gao W, Hillwig ML, Huang L, Cui G, Wang X, Kong J, Yang B, Peters RJ. 2009. A functional genomics approach to tanshinone biosynthesis provides stereochemical insights. Org Lett 11 (22):5170–5173. doi:10.1021/ol902051v.

Ge X, Wu J. 2005. Induction and potentiation of diterpenoid tanshinone accumulation in *Salvia miltiorrhiza* hairy roots by beta-aminobutyric acid. Appl Microbiol Biotechnol 68 (2):183–188. doi:10.1007/s00253-004-1873-2.

Gong Y, Li Y, Abdolmaleky HM, Li L, Zhou JR. 2012. Tanshinones inhibit the growth of breast cancer cells through epigenetic modification of Aurora A expression and function. PLoS One 7 (4):e33656. doi:10.1371/journal.pone.0033656.

Gu XC, Chen JF, Xiao Y, Di P, Xuan HJ, Zhou X, Zhang L, Chen WS. 2012. Overexpression of allene oxide cyclase promoted tanshinone/phenolic acid production in *Salvia miltiorrhiza*. Plant Cell Rep 31 (12):2247–2259. doi:10.1007/s00299-012-1334-9.

Gundlach H, Muller MJ, Kutchan TM, Zenk MH. 1992. Jasmonic acid is a signal transducer in elicitor-induced plant cell cultures. Proc Natl Acad Sci U S A 89 (6):2389–2393.

Guo J, Zhou YJ, Hillwig ML, et al. 2013. CYP76AH1 catalyzes turnover of miltiradiene in tanshinones biosynthesis and enables heterologous production of ferruginol in yeasts. Proc Natl Acad Sci U S A 110 (29):12108–12113. doi:10.1073/pnas. 1218061110.

Hao X, Shi M, Cui L, Xu C, Zhang Y, Kai G. 2015. Effects of methyl jasmonate and salicylic acid on tanshinone production and biosynthetic gene expression in transgenic *Salvia miltiorrhiza* hairy roots. Biotechnol Appl Biochem 62 (1):24–31. doi:10.1002/bab.1236.

Ishiguro S, Nakamura K. 1994. Characterization of a cDNA encoding a novel DNA-binding protein, SPF1, that recognizes SP8 sequences in the 5’ upstream regions of genes coding for sporamin and beta-amylase from sweet potato. Mol Gen Genet 244 (6):563–571.

Jiang W, Fu X, Pan Q, et al. 2016. Overexpression of AaWRKY1 Leads to an Enhanced Content of Artemisinin in *Artemisia annua*. Biomed Res Int 2016:7314971. doi:10.1155/2016/7314971.

Jiang W, Wu J, Zhang Y, Yin L, Lu J. 2015. Isolation of a WRKY30 gene from *Muscadinia rotundifolia (Michx)* and validation of its function under biotic and abiotic stresses. Protoplasma 252 (5):1361–1374. doi:10.1007/s00709-015-0769-6.

Kai G, Xu H, Zhou C, Liao P, Xiao J, Luo X, You L, Zhang L. 2011. Metabolic engineering tanshinone biosynthetic pathway in *Salvia miltiorrhiza* hairy root cultures. Metab Eng 13 (3):319–327. doi:10.1016/j.ymben.2011.02.003.

Kai GY, Liao P, Xu H, Wang J, Zhou CC, Zhou W, Qi YP, Xiao JB, Wang YL, Zhang L. 2012. Molecular mechanism of elicitor-induced tanshinone accumulation in *Salvia miltiorrhiza* hairy root cultures. Acta Physiol Plant 34 (4):1421–1433.

Kai GY, Liao P, Zhang T, Zhou W, Wang J, Xu H, Liu YY, Zhang L. 2010. Characterization, Expression Profiling, and Functional Identification of a Gene Encoding Geranylgeranyl Diphosphate Synthase from *Salvia miltiorrhiza*. Biotechnol Bioproc E 15 (2):236–245.

Kalde M, Barth M, Somssich IE, Lippok B. 2003. Members of the *Arabidopsis* WRKY group III transcription factors are part of different plant defense signaling pathways. Mol Plant Microbe Interact 16 (4):295–305. doi:10.1094/MPMI.2003.16.4.295.

Kato N, Dubouzet E, Kokabu Y, Yoshida S, Taniguchi Y, Dubouzet JG, Yazaki K, Sato F. 2007. Identification of a WRKY protein as a transcriptional regulator of benzylisoquinoline alkaloid biosynthesis in *Coptis japonica*. Plant Cell Physiol 48(1): 8–18. doi:10.1093/pcp/pcl041.

Li JB, Luan YS, Liu Z. 2015. Overexpression of SpWRKY1 promotes resistance to *Phytophthora nicotianae* and tolerance to salt and drought stress in transgenic tobacco. Physiol Plant 155 (3):248–266. doi:10.1111/ppl.12315.

Liao P, Zhou W, Zhang L, Wang J, Yan XM, Zhang Y, Zhang R, Li L, Zhou GY, Kai GY. 2009 Molecular cloning, characterization and expression analysis of a new gene encoding 3-hydroxy-3-methylglutaryl coenzyme A reductase from *Salvia miltiorrhiza*. Acta Physiol Plant 31 (3):565–572.

Liu Q, Liu Y, Tang Y, Chen J, Ding W. 2017. Overexpression of NtWRKY50 Increases Resistance to *Ralstonia solanacearum* and Alters Salicylic Acid and Jasmonic Acid Production in Tobacco. Front Plant Sci 8:1710. doi:10.3389/fpls.2017.01710.

Liu X, Song Y, Xing F, Wang N, Wen F, Zhu C. 2016. GhWRKY25, a group I WRKY gene from cotton, confers differential tolerance to abiotic and biotic stresses in transgenic *Nicotiana benthamiana*. Protoplasma 253 (5):1265–1281. doi:10.1007/s00709-015-0885-3.

Lu M, Wang LF, Du XH, Yu YK, Pan JB, Nan ZJ, Han J, Wang WX, Zhang QZ, Sun QP. 2015. Molecular cloning and expression analysis of jasmonic acid dependent but salicylic acid independent LeWRKY1. Genetics and Molecular Research 14(4):15390–15398.

Ma D, Pu G, Lei C, et al. 2009. Isolation and characterization of AaWRKY1, an *Artemisia annua* transcription factor that regulates the amorpha-4,11-diene synthase gene, a key gene of artemisinin biosynthesis. Plant Cell Physiol 50 (12):2146–2161. doi:10.1093/pcp/pcp149.

Ma S, Gong Q, Bohnert HJ. 2006. Dissecting salt stress pathways. J Exp Bot 57 (5): 1097–1107. doi:10.1093/jxb/erj098.

Ma Y, Yuan L, Wu B, Li X, Chen S, Lu S. 2012. Genome-wide identification and characterization of novel genes involved in terpenoid biosynthesis in *Salvia miltiorrhiza*. J Exp Bot 63 (7):2809–2823. doi:10.1093/jxb/err466.

Phukan UJ, Jeena GS, Shukla RK. 2016. WRKY Transcription Factors: Molecular Regulation and Stress Responses in Plants. Front Plant Sci 7:760. doi:10.3389/fpls.2016.00760.

Qiu D, Xiao J, Ding X, Xiong M, Cai M, Cao Y, Li X, Xu C, Wang S. 2007. OsWRKY13 mediates rice disease resistance by regulating defense-related genes in salicylate- and jasmonate-dependent signaling. Mol Plant Microbe Interact 20(5): 492–499. doi:10.1094/MPMI-20-5-0492.

Rushton PJ, Somssich IE, Ringler P, Shen QJ. 2010. WRKY transcription factors. Trends Plant Sci 15 (5):247–258. doi:10.1016/j.tplants.2010.02.006.

Rushton PJ, Torres JT, Parniske M, Wernert P, Hahlbrock K, Somssich IE. 1996. Interaction of elicitor-induced DNA-binding proteins with elicitor response elements in the promoters of parsley PR1 genes. EMBO J 15 (20):5690–5700.

Shi M, Luo X, Ju G, Li L, Huang S, Zhang T, Wang H, Kai G. 2016a. Enhanced Diterpene Tanshinone Accumulation and Bioactivity of Transgenic *Salvia miltiorrhiza* Hairy Roots by Pathway Engineering. J Agric Food Chem 64 (12):2523–2530. doi:10.1021 /acs jafc.5b04697.

Shi M, Luo X, Ju G, Yu X, Hao X, Huang Q, Xiao J, Cui L, Kai G. 2014. Increased accumulation of the cardio-cerebrovascular disease treatment drug tanshinone in *Salvia miltiorrhiza* hairy roots by the enzymes 3-hydroxy-3-methylglutaryl CoA reductase and 1-deoxy-D-xylulose 5-phosphate reductoisomerase. Funct Integr Genomics 14 (3):603–615. doi:10.1007/s 10142-014-0385-0.

Shi M, Zhou W, Zhang J, Huang S, Wang H, Kai G. 2016b. Methyl jasmonate induction of tanshinone biosynthesis in *Salvia miltiorrhiza* hairy roots is mediated by JASMONATE ZIM-DOMAIN repressor proteins. Sci Rep 6:20919. doi:10.1038/srep20919.

Singh AK, Kumar SR, Dwivedi V, Rai A, Pal S, Shasany AK, Nagegowda DA. 2017. A WRKY transcription factor from *Withania somnifera* regulates triterpenoid withanolide accumulation and biotic stress tolerance through modulation of phytosterol and defensepathways. New Phytol 215 (3): 1115–1131. doi:10.1111/nph.14663.

Suttipanta N, Pattanaik S, Kulshrestha M, Patra B, Singh SK, Yuan L. 2011. The transcription factor CrWRKY1 positively regulates the terpenoid indole alkaloid biosynthesis in *Catharanthus roseus*. Plant Physiol 157 (4): 2081–2093. doi:10.1104/pp.111.181834.

Tsuda K, Sato M, Stoddard T, Glazebrook J, Katagiri F. 2009. Network properties of robust immunity in plants. PLoS Genet 5 (12):e1000772. doi:10.1371/journal.pgen.1000772.

Wang F, Chen HW, Li QT, et al. 2015. GmWRKY27 interacts with GmMYB174 to reduce expression of GmNAC29 for stress tolerance in soybean plants. Plant J 83(2): 224–236. doi:10.1111/tpj.12879.

Wang Q, Wang M, Zhang X, Hao B, Kaushik SK, Pan Y. 2011. WRKY gene family evolution in *Arabidopsis thaliana*. Genetica 139 (8):973–983. doi:10.1007/s10709-011-9599-4.

Wu KL, Guo ZJ, Wang HH, Li J. 2005. The WRKY family of transcription factors in rice and *Arabidopsis* and their origins. DNA Res 12 (1):9–26.

Xie Z, Zhang ZL, Zou X, Huang J, Ruas P, Thompson D, Shen QJ. 2005. Annotations and functional analyses of the rice WRKY gene superfamily reveal positive and negative regulators of abscisic acid signaling in aleurone cells. Plant Physiol 137 (1):176–189. doi:10.1104/pp.104.054312.

Xu H, Zhang L, Zhou CC, Xiao JB, Liao P, Kai GY. 2010. Metabolic regulation and genetic engineering of pharmaceutical component tanshinone biosynthesis in *Salvia miltiorrhiza*. J Med Plants Res 4 (24):2591–2597.

Xu M, Hao H, Jiang L, Long F, Wei Y, Ji H, Sun B, Peng Y, Wang G, Ju W, Li P. 2015. In vitro inhibitory effects of ethanol extract of Danshen *(Salvia miltiorrhiza)* and its components on the catalytic activity of soluble epoxide hydrolase. Phytomedicine 22 (4):444–451. doi:10.1016/j.phymed.2015.02.001.

Xu YH, Wang JW, Wang S, Wang JY, Chen XY. 2004. Characterization of GaWRKY1, a cotton transcription factor that regulates the sesquiterpene synthase gene (+)-delta-cadinene synthase-A. Plant Physiol 135 (1):507–515. doi:10.1104/pp.104.038612.

Yan XM, Zhang L, Wang J, Liao P, Zhang Y, Zhang R, Kai GY. 2009. Molecular characterization and expression of 1-deoxy-d-xylulose 5-phosphate reductoisomerase (DXR) gene from *Salvia miltiorrhiza*. Acta Physiol Plant 31 (5):1015–1022.

Yin G, Xu H, Xiao S, Qin Y, Li Y, Yan Y, Hu Y. 2013. The large soybean (Glycine max) WRKY TF family expanded by segmental duplication events and subsequent divergent selection among subgroups. BMC Plant Biol 13:148. doi:10.1186/1471-2229-13-148.

Yoda H, Ogawa M, Yamaguchi Y, Koizumi N, Kusano T, Sano H. 2002. Identification of early-responsive genes associated with the hypersensitive response to tobacco mosaic virus and characterization of a WRKY-type transcription factor in tobacco plants. Mol Genet Genomics 267 (2): 154–161. doi: 10.1007/s00438-002-0651-z.

Zhang G, Tian Y, Zhang J, Shu L, Yang S, Wang W, Sheng J, Dong Y, Chen W. 2015. Hybrid de novo genome assembly of the Chinese herbal plant danshen *(Salvia miltiorrhiza Bunge)*. Gigascience 4:62. doi:10.1186/s13742-015-0104-3.

Zhang J, Zhou L, Zheng X, Zhang J, Yang L, Tan R, Zhao S. 2017. Overexpression of SmMYB9b enhances tanshinone concentration in *Salvia miltiorrhiza* hairy roots. Plant Cell Rep 36 (8): 1297–1309. doi:10.1007/s00299-017-2154-8.

Zhang L, Yan XM, Wang J, Li SS, Liao P, Kai GY. 2011. Molecular cloning and expression analysis of a new putative gene encoding 3-hydroxy-3-methylglutaryl-CoA synthase from *Salvia miltiorrhiza*. Acta Physiol Plant 33 (3):953–961.

Zhang Y, Li X, Wang Z. 2010. Antioxidant activities of leaf extract of *Salvia miltiorrhiza Bunge* and related phenolic constituents. Food Chem Toxicol 48 (10):2656–2662. doi:10.1016/j.fct.2010.06.036.

Zhang Y, Wang L. 2005. The WRKY transcription factor superfamily: its origin in eukaryotes and expansion in plants. BMC Evol Biol 5:1. doi:10.1186/1471-2148-5-1.

Zhao S, Zhang J, Tan R, Yang L, Zheng X. 2015. Enhancing diterpenoid concentration in *Salvia miltiorrhiza* hairy roots through pathway engineering with maize C1 transcription factor. J Exp Bot 66 (22):7211–7226. doi:10.1093/jxb/erv418.

Zhou W, Jiang YR, Zhang W, Xu GF, Rong JK. 2013. Characterization of Large Chromosome Segment Introgressions from *Triticum turgidum subsp* dicoccoides into Bread Wheat with Simple Sequence Repeat Markers. Crop Sci 53 (4):1555–1565.

Zhou W, Huang F, Li S, Wang Y, Zhou C, Shi M, Wang J, Chen Y, Wang Y, Wang H, Kai G. 2016a. Molecular cloning and characterization of two 1-deoxy-d-xylulose-5-phosphate synthase genes involved in tanshinone biosynthesis in *Salvia miltiorrhiza*. MolBreeding 36(9):124, DOI:10.1007/s11032-016-0550-3.

Zhou Y, Sun W, Chen J, et al. 2016b. SmMYC2a and SmMYC2b played similar but irreplaceable roles in regulating the biosynthesis of tanshinones and phenolic acids in *Salvia miltiorrhiza*. Sci Rep 6:22852. doi:10.1038/srep22852.

Zhou W, Wu S, Ding M, Li J, Shi Z, Wei W, Guo JL, Zhang H, Jiang YR, Rong JK. 2016c. Mapping of Ppd-B1, a major candidate gene for late heading on wild emmer chromosome arm 2bs and assessment of its interactions with early heading QTLs on 3AL. PLoS ONE 11(2): e0147377. doi:10.1371/journal.pone.0147377.

Zhou W, Huang Q, Wu X, Zhou Z, Ding M, Shi M, Huang F, Li S, Wang Y, Kai G. 2017. Comprehensive transcriptome profiling of *Salvia miltiorrhiza* for discovery of genes associated with the biosynthesis of tanshinones and phenolic acids. Sci Rep 7 (1):10554. doi:10.1038/s41598-017-10215-2.

